# Bacterial Controller Aided Wound Healing: A Case Study in Dynamical Population Controller Design

**DOI:** 10.1101/659714

**Authors:** Leopold N. Green, Chelsea Y. Hu, Xinying Ren, Richard M. Murray

## Abstract

Wound healing is a complicated biological process consisting of many types of cellular dynamics and functions regulated by chemical and molecular signals. Recent advances in synthetic biology have made it possible to predictably design and build closed-loop controllers that can function appropriately alongside biological species. In this paper we develop a simple dynamical population model mimicking the sequential relay-like dynamics of cellular populations involved in the wound healing process. Our model consists of four nodes and five signals whose parameters we can tune to simulate various chronic healing conditions. We also develop a set of regulator functions based on type-1 incoherent feed forward loops (IFFL) that can sense the change from acute healing to incomplete chronic wounds, improving the system in a timely manner. Both the wound healing and type-1 IFFL controller architectures are compatible with available synthetic biology experimental tools for potential applications.

## Introduction

Wound healing is a dynamical, multi-cellular process regulated by a complicated network of propagating cell signals [1]. A healthy response to tissue injury relies on a systematic cascade of events known as acute wound healing. Acute healing is classically defined by four consecutive phases distinct in function and histological characteristics: the hemostasis phase involving blood coagulation, the inflammatory phase in which the wound is debrided of foreign material, the proliferation phase when granulation tissue forms and the wound closes, and finally the remodelling phase which includes improving the tensile strength of the wound [2]. Together, the phases demonstrate relay-like dynamics of cellular densities and functions, coordinated by the secretion of various signals [3, 4].

Some of the more prominent cells involved in wound healing are platelets, neutrophils, macrophages, fibroblasts, and endothelial cells. The sequential flow of varying cell populations into the wound, illustrated in Figure 1A, is carried out by a combination of cell migration, infiltration, proliferation, and differentiation. It is also controlled by an equally sophisticated signaling network of growth factors, cytokines, and chemokines [5, 6]. Platelets are the first cell type to enter the wound, beginning at the moment of injury. Platelets promote the formation of blood clots, activate coagulation, and recruit various inflammatory cells into the wound by releasing pro-inflammatory growth factors [7] and cytokines.

**Figure 1:**
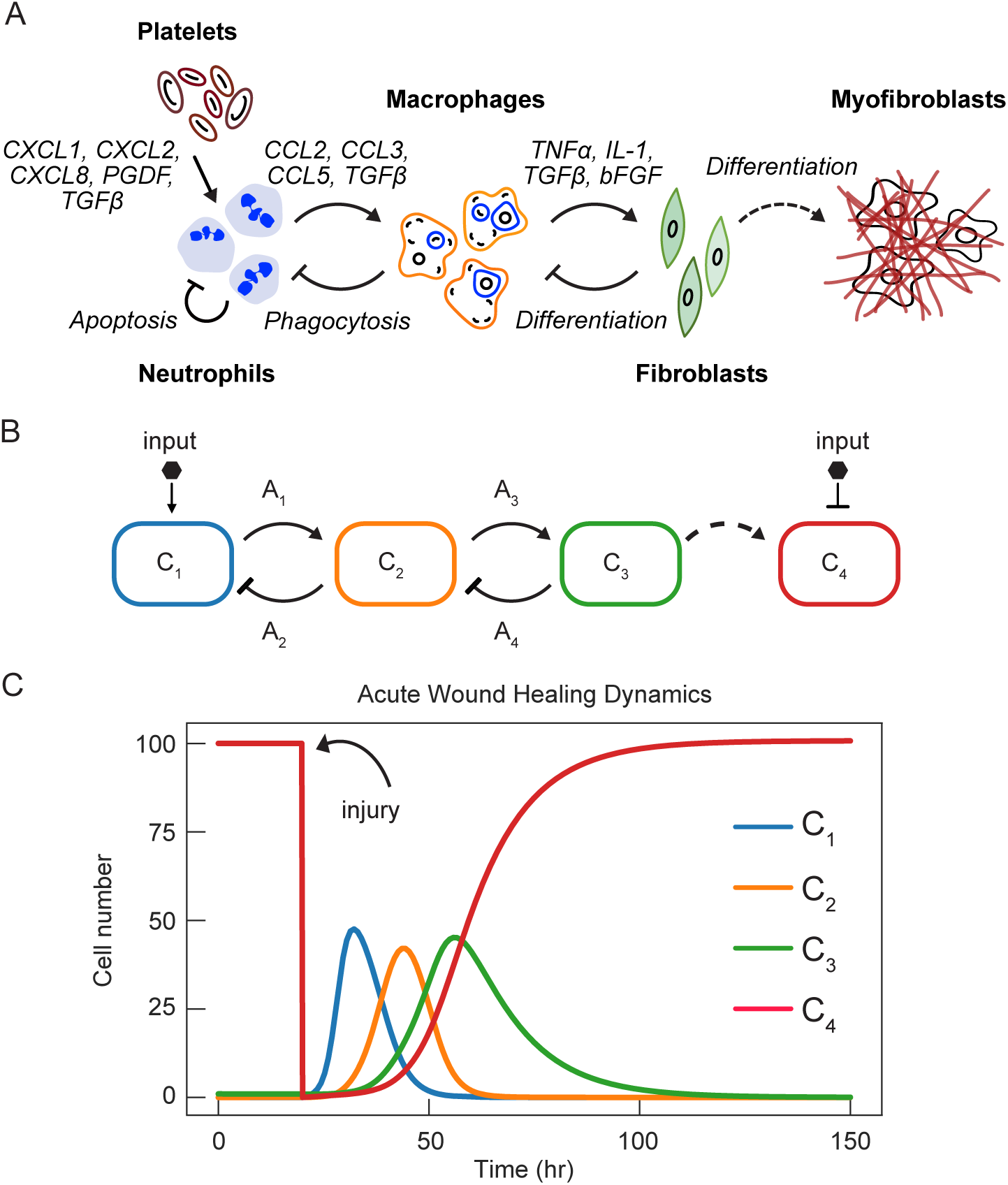
Simulating the wound healing dynamics with a reduced model. A. Diagram of wound healing signal and cell propagation dynamics. Upon injury the hemostasis stage occurs, preventing further blood loss via platelet activation. Platelets release chemokines and other growth factors, recruiting neutrophils into the wound. Neutrophils are the first inflammatory cell type to enter the wound. Mature neutrophils undergo apoptosis, releasing signals to recruit macrophages. Macrophages continue the debridement process of the wound consuming non-active neutrophils and release signals to promote the migration of fibroblasts. The presence of fibroblasts corresponds with the proliferation phase. As the extra-cellular matrix (ECM) is reconstructed, various proteins and chemical signals released by the ECM activate macrophage differentiation from M1 to M2, as well as the differentiation of fibroblasts to myofibroblasts. B. Circuit diagram used to demonstrate acute wound healing computationally. *C*_1_ = Neutrophils; *C*_2_ = Macrophages; *C*_3_ = Fibroblasts; *C*_4_ = Myofibroblasts/Collagen. *A*_1_ and *A*_3_ are signals for growth; *A*_2_ and *A*_4_ are signals for death. C. Simulation of acute wound healing dynamics using parameters listed in Table 2 and ODEs in equations (1-8).

As the first circulating inflammatory cells to enter the site of injury, neutrophils overtake platelets as the predominant cell in the wound, protecting the host from pathogenic infections. Neutrophils secrete antimicrobial proteins that help degrade potential pathogens and produce proinflammatory signals for self-proliferation and macrophage migration into the wound [8]. Macrophages continue the cleaning of the wound by microbial phagocytosis and the digestion of cellular debris. As more monocytes differentiate into macrophages, the once dominant neutrophil populations decrease due to pro-apoptotic endogenous signals of mature neutrophils [9, 10, 11]. By the time the inflammatory cycle ends, the macrophage-derived growth factors reach the optimal level causing an influx of fibroblasts into the wound [12, 13].

During the proliferation phase, fibroblast and epithelial cells are the most prominent cell types present. As fibroblasts continue to migrate into the wound, components of the extra-cellular matrix (ECM) including collagen, are produced, promoting wound closure and increasing tensile strength [14]. Following proliferation and ECM formation, healing enters the remodelling phase. In some cases, the remodelling phase can last many years as the collagen and granulation tissues are constantly being reorganized. Also during the remodelling phase, fibroblasts differentiate into contractile myofibroblasts resulting in minimal scar tissue and preserved tissue function [15].

Even in the simplified description above, it is easy to appreciate the complexity and coordination of the healing process. When deviations from the acute process occurs, healing is delayed, and ften times chronic, non-healing wounds persist. Chronic wounds are often difficult to treat due to the large array of molecular and cellular pathology within the chronic environment [16, 17]. Effective therapeutics must be coordinated temporally and molecularly to regulate altered signals appropriately.

Here, we present a multi-layer control strategy for an engineered healing regulator system. The first layer is designed to mimic cellular population dynamics of acute healing with tunable parameters for induced chronic wound healing. The second layer regulates against chronic conditions via pulse signals from activated cell types. We show that a type-1 pulse generating incoherent feed-forward loop (IFFL) architecture is an effective design choice for our desired functions.

## Wound healing design strategy

Our proposed wound healing circuit is an abstraction of physiological wound healing focusing on only a few of many well studied cell types of known importance and characterized dynamics: neutrophils (*C*_1_), macrophages (*C*_2_), fibroblasts (*C*_3_) and myofibroblasts (*C*_4_), illustrated in Figure 1A and B. The dynamics of the skin healing cascade depend heavily on the population of each cell type and the major chemical mediators developed in each phase. From the proposed circuit diagram in Figure 1, we developed a deterministic model of two coupled negative feedback modules producing sequential pulses of signals resembling the physiological acute healing process dynamics.

We analyzed the progression of healing by the population dynamics of each of the four cell types over time. Our goal is to eventually extend this computational model to an experimental demonstration of wound healing using engineered *E. coli* as our cellular chassis. Therefore, our design strategies are based on bacterial implementation and dynamics. Table 1 summarizes the species involved in the mechanisms described in the subsequent sections. Table 2 summarizes the parameters used to simulate acute wound healing dynamics in a bacterial system.

**Table 1:** Summary of species in our wound healing model

| Species | Description |
| --- | --- |
| $C_1$ | Population of $C_1$ |
| $C_2$ | Population of $C_2$ |
| $C_3$ | population of $C_3$ |
| $C_4$ | Population of $C_4$ |
| $A_1$ | AHL signal produced in $C_1$ |
| $A_2$ | AHL signal produced in $C_2$ |
| $A_3$ | AHL signal produced in $C_2$ |
| $A_4$ | AHL signal produced in $C_3$ |

### Acute wound healing dynamics

To simplify the model, cell proliferation, migration (in), and infiltration are modeled as cell growth; while apoptosis, phagocytosis, cell migration (out), and, in the case of macrophage dynamics, cell differentiation are modeled as cell death. All communication signals (species *A*_1_, *A*_2_, *A*_3_ and *A*_4_) are modeled as quorum sensing molecules that permeate through cell walls instantly. The topology of the circuit design is illustrated in Figure 1B.

The minimal model for our wound healing dynamics simulations, are based on the following assumptions:

- Every cell in the population of a given cell type contains identical circuit function; the activation of either cell growth or cell death is triggered by two orthogonal signals.
- Cell growth has logistic kinetics with a growth rate constant of 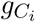 and a carrying capacity of *C*_*max*_. The growth rate of *C*_*i*_ is proportional to the total cell population, where *i* = 1, 2, 3, 4.
- Activation of both cell growth and cell death of species *C*_*i*_ by signal *A*_*j*_ is governed by a first order Hill function with a dissociation constant 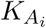, where *j* = 1, 2, 3, 4, and *i* = 1, 2, 3, 4.
- The dilution/basal death rate of cells is much slower than the kinetics of induced death rate in the circuit; thus dilution/basal death rates are negligible. Lysis induced death of *C*_1_ and *C*_2_ is activated by *A*_2_ and *A*_4_ respectively.
- The production of signal species *A*_*j*_ is characterized by its maximal rate of 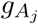.
- The dilution/degradation rate of signal species *A*_*j*_ is described as 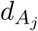.

We obtain the following model for all acute wound healing cellular species *C*_1_, *C*_2_, *C*_3_, and *C*_4_, as well as signaling species *A*_1_, *A*_2_, *A*_3_ and *A*_4_. The first term in equations (1-3) represents recruiting *C*_1_, *C*_2_, and *C*_3_ with signals *µ, A*_1_, and *A*_3_ respectively. Once cells are present, recruitment stops, and signal induced cell growth is activated.

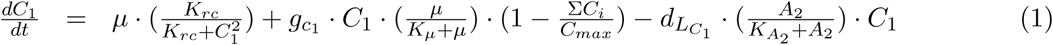

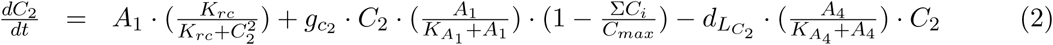

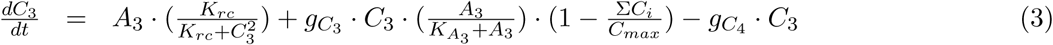

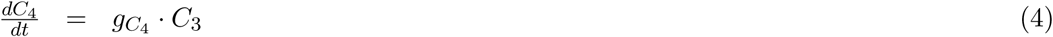

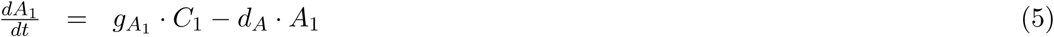

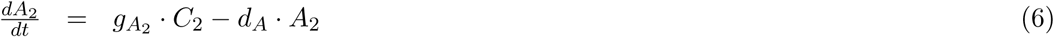

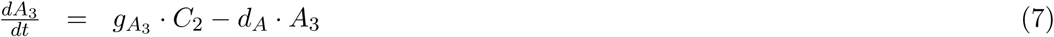

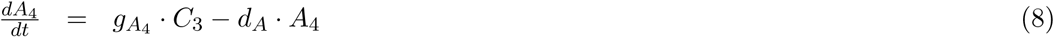

As mentioned earlier, the first layer of the control systems is a simulation of cellular population dynamics that represents that of physiological acute healing. In our simulation, species *C*_4_ represents complete wound contraction (healthy). Upon injury, via activation of input signal *µ*, we simulate breaching of the skin barrier (setting initial condition of *C*_4_ to zero), while simultaneously inducing the growth of species *C*_1_ (initiation of inflammation phase) as shown in equation (1). The first term in equation (1) describes the recruitment of *C*_1_ into the system. This recruitment is turned off as soon as a small amount of *C*_1_ is present. After the recruitment, the species *µ* regulates the growth of *C*_1_ at the maximum rate of 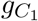.

As the *C*_1_ population increases, the constitutive expression of signaling molecule *A*_1_ accumulates both in the cell and globally at a rate of 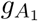 as described in equation (5). We assume permeation of the signal is instantaneous, therefore the concentration of the signal inside and outside of the cell is the same. Species *A*_1_ activates the growth of *C*_2_ (similar of the mechanism of *µ* activating *C*_1_), equation (2). Cell species *C*_2_ constitutively produces two signals, *A*_2_ in equation (6) and signal *A*_3_ in equation (7). While *A*_2_ negatively regulates *C*_1_ by activating its death at rate 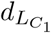, *A*_3_ progresses healing to the next phase by recruiting *C*_3_, equation (3).

In physiological wound healing, species *C*_2_ (macrophage) serves two purposes: (1) finishing the debridement function of the inflammatory phase (lowering concentration of *C*_1_), and (2) activating anti-inflammatory signals for the recruitment of fibroblasts *C*_3_ (proliferation phase). Once activated, *C*_3_ produces a signal *A*_4_ that negatively regulates *C*_2_ by activating the death of *C*_2_. With species *C*_1_ no longer in the system, and species *C*_2_ approaching its depletion, *C*_3_ becomes the dominant cell type in the wound. Physiologically, at this stage, the fibroblast is responsible for establishing wound closure by secreting collagen.

Once collagen is placed in the wound, the final healing stage is activated. At this stage, the wound is free of any damaged cellular debris and pathogens from injury through the *C*_1_ and *C*_2_ phases, and has a strong, functional matrix of collagen. The final stage of wound healing, wound contraction, is governed by the differentiation of fibroblasts into myofibroblasts, modeled as *C*_3_ converting to *C*_4_ as shown in equation (4). Based on parameters listed in Table 2, the acute wound healing model demonstrates the similar sequential relay dynamics found in physiological wound healing (Figure 1C).

**Table 2:** Summary of parameters in our would healing model

| Parameters | Description | Value |
| --- | --- | --- |
| $g_{C_1}$ | Cell growth rate of $C_1$ | $1.5 \text{ h}^{-1}$ |
| $g_{C_2}$ | Cell growth rate of $C_2$ | $1.5 \text{ h}^{-1}$ |
| $g_{C_3}$ | Cell growth rate of $C_3$ | $2.8 \text{ h}^{-1}$ |
| $g_{C_4}$ | Rate of differentiation of $C_3$ to $C_4$ | $0.08 \text{ h}^{-1}$ |
| $g_{A_1}$ | Rate of $A_1$ production in $C_1$ | $5 \text{ nM h}^{-1}$ |
| $g_{A_2}$ | Rate of $A_2$ production in $C_2$ | $5 \text{ nM h}^{-1}$ |
| $g_{A_3}$ | Rate of $A_3$ production in $C_3$ | $5 \text{ nM h}^{-1}$ |
| $g_{A_4}$ | Rate of $A_4$ production in $C_4$ | $5 \text{ nM h}^{-1}$ |
| $C_{max}$ | Maximal cell density | 100 cells |
| $K_\mu$ | Activating/repression constant of $\mu$ | 1 nM |
| $K_{A_1}$ | Activating/repression constant of $A_1$ | 200 nM |
| $K_{A_2}$ | Activating/repression constant of $A_2$ | 200 nM |
| $K_{A_3}$ | Activating/repression constant of $A_3$ | 400 nM |
| $K_{A_4}$ | Activating/repression constant of $A_4$ | 400 nM |
| $K_{rc}$ | Species recruiting constant | 0.1 nM |
| $d_{C_R}$ | Natural cell death rate of $R_1$ and $R_2$ | $0.1 \text{ h}^{-1}$ |
| $d_{L_{C_1}}$ | Maximal $C_1$ death rate due to lysis | $0.6 \text{ h}^{-1}$ |
| $d_{L_{C_2}}$ | Maximal $C_2$ death rate due to lysis | $0.6 \text{ h}^{-1}$ |
| $d_A$ | Deg. rate of signaling molecules | $0.5 \text{ h}^{-1}$ |
| $\mu$ | Inducer signal species initiating propagation | 0 or 1 |

**Table 3:**
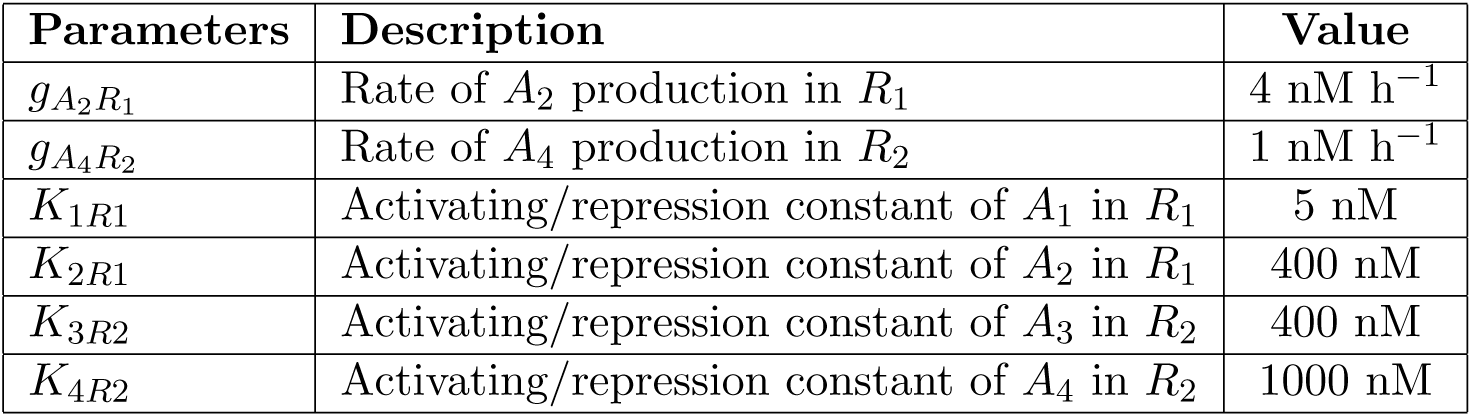
Parameters for the regulator cells

We further analyzed the cellular dynamics of our first layer acute healing circuit design. In Figure 2, the introduction of population disturbances were simulated by decreasing the cellular concentrations of *C*_1_ through *C*_4_ during their respective phase of healing. Perturbing cellular species resulted in minimal delay in healing and alterations to cellular dynamics of each of the cell types in the system. Notice the perturbations of *C*_2_ in Figure 2B slightly delay healing. This is because species *C*_2_ is responsible for activating the death of *C*_1_, as well as the growth of *C*_3_. Perturbing the population of *C*_2_ allows for the prolonged presence of *C*_1_, delaying overall healing. A perturbations of *C*_4_ when approaching its peak density (complete healing), simulate a secondary injury within the original wounded region, triggering the cascading dynamics of healing starting from the first stage, growth of *C*_1_ (Figure 2D).

**Figure 2:**
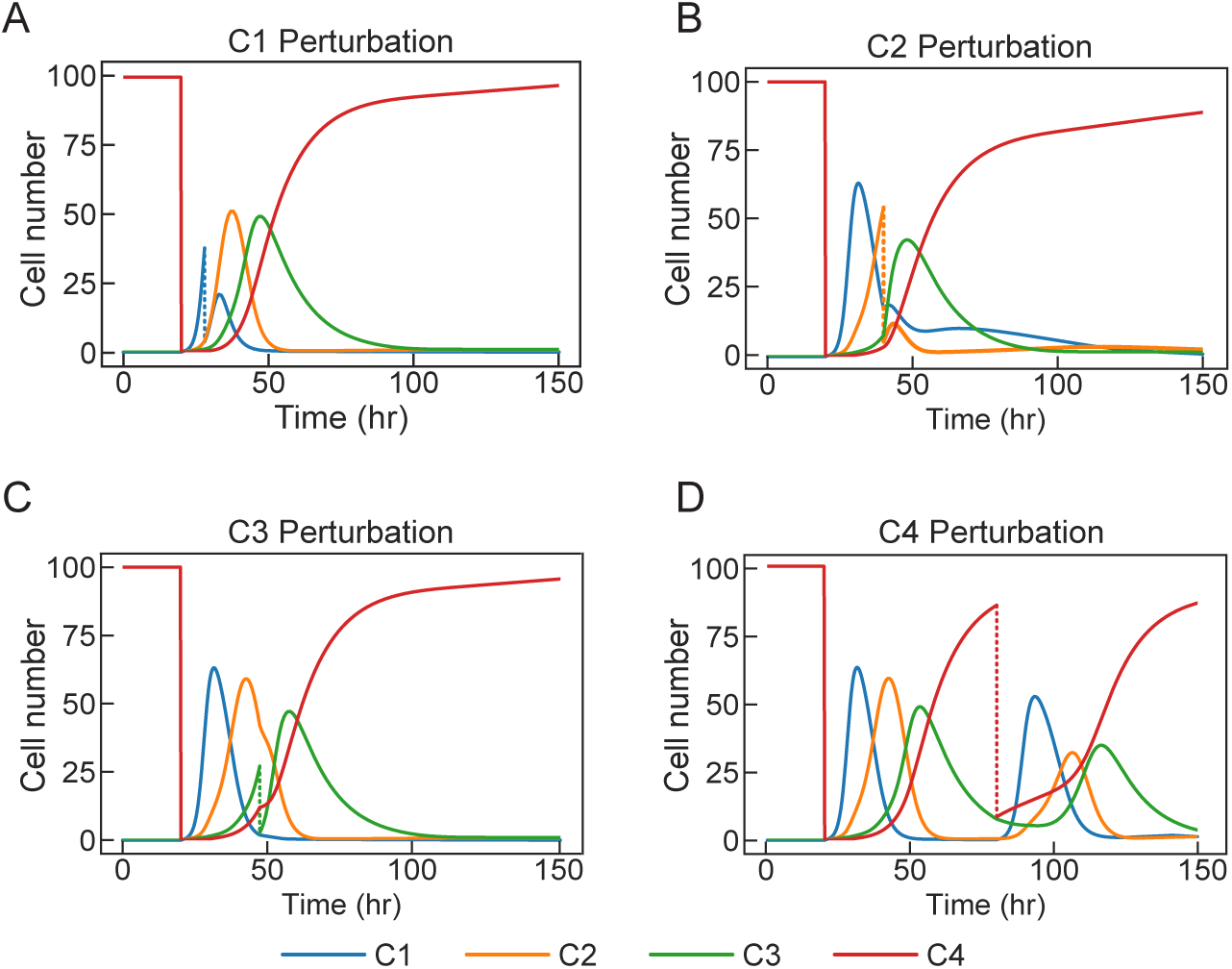
Cellular density perturbation in acute wound healing. Perturbations resulting in the removal of each cell type at their respective times: A. depletion of species *C*_1_ at t = 28 hr, B. depletion of species *C*_2_ at t = 40 hr, C. depletion of species *C*_3_ at t = 47 hr, and D. depletion of species *C*_4_ at t = 80 hr. 6

### Chronic wound healing conditions

Whether caused by dermal injury or surgical procedure, acute wound healing progresses steadily and predictably through all of the healing stages. Ultimately, the time span and outcome of acute healing will depend on the wound’s location, size, depth, and trauma type. Chronic wounds, on the other hand, are defined as wounds that have not fully healed, normally surpassing 30 days of recovery. Many factors including oxygenation, infection, age, stress, diabetes, medications, and nutrition, are known to interfere with the wound’s ability to progress through normal healing [18, 19].

The orchestrated interactions of various cell types, extra-cellular components, growth factors, and cytokines together play important roles in each of the different stages during healing [20]. Therefore, an imbalance to any of these elements may lead to either prolonged healing, or excessive scarring. Most commonly, chronic wounds are thought to be stalled in either the inflammatory phase or the proliferative phase [21, 22]. There are many factors in physiological wound healing that can go awry, leading to non-healing wounds. In this section we focus on four conditions that lead to hyper-inflammation. In Figure 3, we analyze the sequential population dynamics of chronic wound conditions based on the following scenarios:

**Figure 3:**
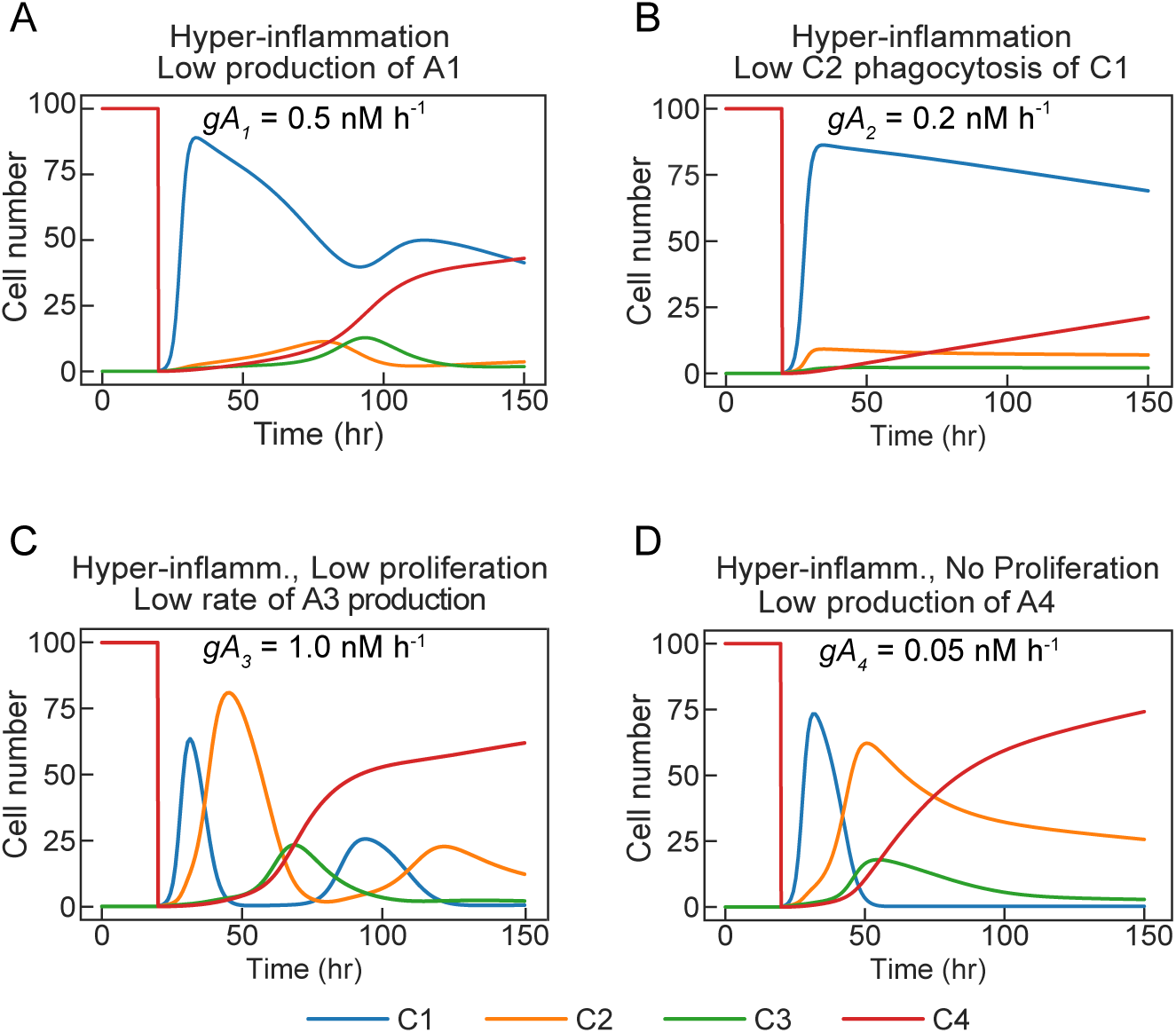
Simulations of chronic wound healing dynamics. A. Low production rate of signal 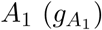, which recruits *C*_2_. B. Low production rate of signal 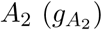, which induces lysis expression in *C*_1_. C. Low production rate of signal 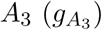, which induces proliferation (growth) of cell *C*_3_ D. Low production rate of signal 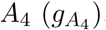, which induces lysis expression in *C*_2_.

- Figure 3A: Hyper-inflammation condition caused by low *A*_1_ signal production 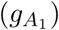 in cell *C*_1_. In this system, we have prolonged activation of *C*_1_, which does not properly recruit *C*_2_ into the system. Without *C*_2_, the wound will remain stuck in the hyper-inflammatory state. In a healthy wound environment, neutrophils recruit the second inflammatory cell species macrophages (*C*_2_), for healthy healing progression. Macrophages play an important role in wound debridement as well as the initiation of the proliferation stage.
- Figure 3B: Hyper-inflammation condition caused by low *A*_2_ signal production 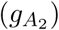 in cell *C*_2_. Similar to the results of Figure 3A, low *A*_2_ production allows for species *C*_1_ to persist in the wound. The action of macrophage phagocytosis of neutrophils (*C*_1_) is modeled as inducible lysis gene expression. The extended presence of neutrophils may result in increased levels of pro-inflammatory signals, resulting in chronic wound healing.
- Figure 3C: Hyper-inflammation conditions caused by a low production rate of signal *A*_3_ in cell *C*_2_. This simulates low proliferation of fibroblasts or a slow transition rate from the inflammatory phase to the proliferation phase. Signal *A*_3_ is used to activated the growth of species *C*_3_. When the rate of *A*_3_ signal production is low, the wound gets stuck in oscillatory dynamics between *C*_1_ and *C*_2_. Physiologically, an improper transition from the inflammatory phase to the proliferation phase could result in increased tissue damage and increased infection rate.
- Figure 3D: Chronic wound healing dynamics of an impaired inflammation to proliferation, simulated with a slow production rate of signal *A*_4_ in cell *C*_3_. Recruited fibroblasts initiate the reconstruction phase in the wound by secreting many ECM signals. ECM signals along with the influx of fibroblasts into the injury site, are responsible for the transition of proinflammatory macrophage cells M1 into anti-inflammatory macrophage cells M2. In cases where the fibroblast function is low, M1 macrophages persist reverting the wound back into a chronic inflammatory condition.

## Regulator cells design strategy

As described earlier, our wound healing circuit consists of two negative feedback modules sharing a common cell type, species *C*_2_: (1) *C*_1_ activating *C*_2_ growth while *C*_2_ represses *C*_1_ (activating death) and (2) *C*_2_ activating *C*_3_ growth while *C*_3_ represses *C*_2_ (also activating death). These interactions are shown in Figure 4A.

**Figure 4:**
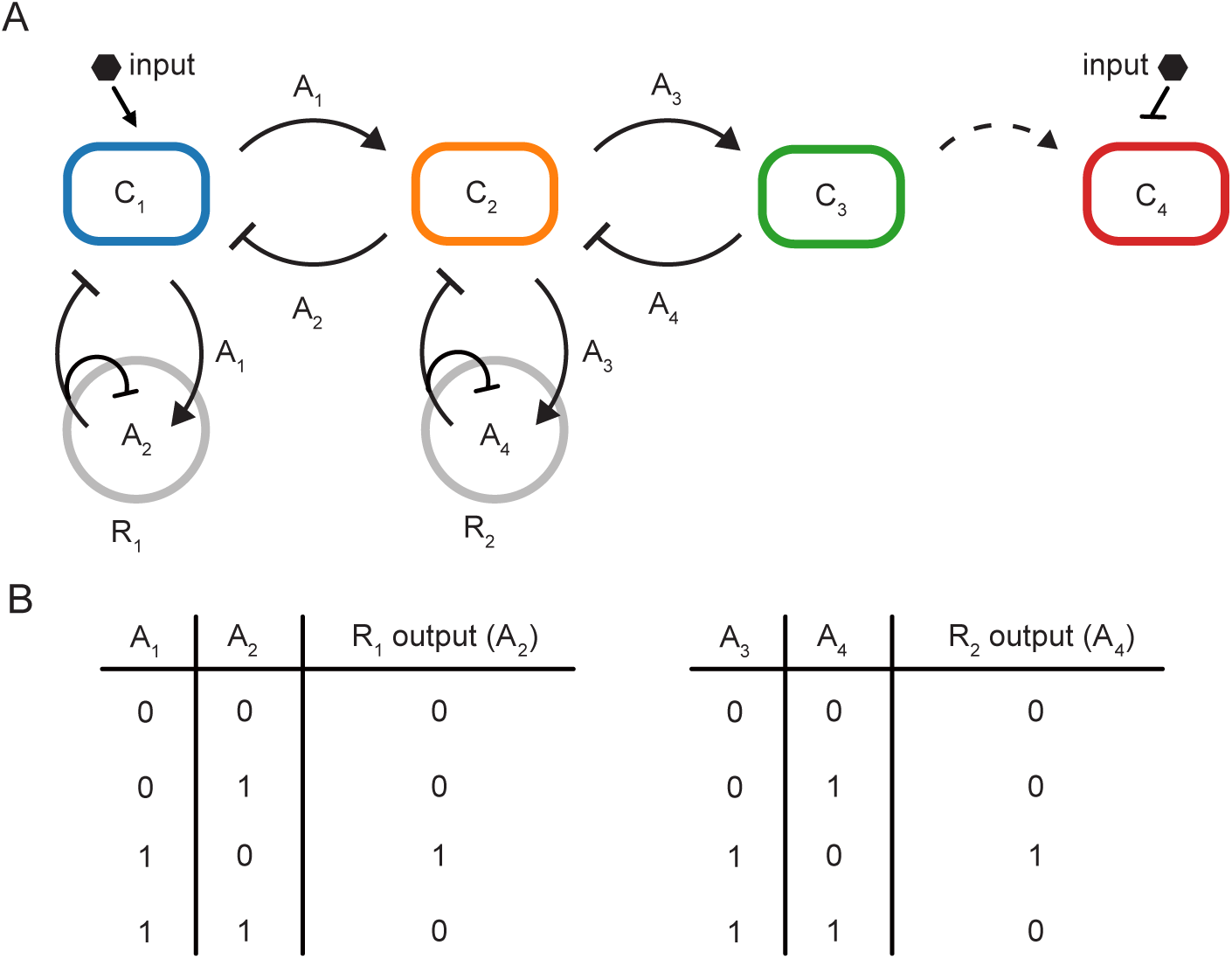
Regulator design: Two coupled incoherent feed-forward loop circuits. A. Circuit diagram of layer 1 wound healing circuit, coupled with layer 2 circuit that consists of two pulse generating regulator cells *R*_1_ and *R*_2_. B. Truth table describing the logic of the regulator cells producing output species *A*_2_ and *A*_4_ in response to the environmental signals.

Here we designed regulator cells, keeping in mind the physiological rules of timely and functional acute wound healing. We propose two coupled type–1 IFFL regulator cells to produce signaling pulses, illustrated in Figure 4 under desired conditions [23, 24]. Species *R*_1_ receives input *A*_1_ and produces output *A*_2_. When signal *A*_2_ is low, signal *A*_1_ is secreted from wound healing species *C*_1_. A high concentration of *C*_1_ activates the production of *A*_2_ in the wound environment, promoting the death of *C*_1_. In summary, high *A*_1_ and low *A*_2_ conditions results in pulsed *A*_2_ production in *R*_1_, preventing the persistence of high *C*_1_ hyper-inflammation. Due to the negative auto-regulation of *A*_2_ in *R*_1_, the production of *A*_2_ from *R*_1_ is turned off as soon as *A*_2_ becomes abundant enough to prevent further potential interference between controller cells and the wound tissues. The second controller *R*_2_ is identical to *R*_1_, with signal *A*_3_ as the input signal and *A*_4_ as the output signal. Figure 4A illustrates the detailed interaction between the wound dynamics and the regulator cells where equation (6) and equation (8) in the acute wound healing model are updated into equation (9) and equation (10), respectively.

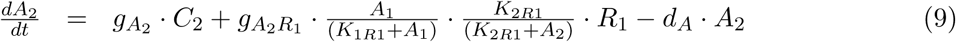

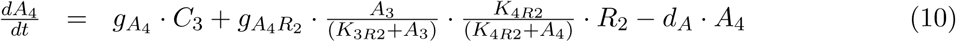

Using the chronic wound condition parameters, we analyzed the effectiveness of our regulator controller cells against selected chronic conditions. Simulations of chronic healing dynamics in the presence of regulator cells shown in Figure 5 demonstrate the controller’s ability to improve the altered cellular dynamics associated with chronic healing. Our coupled pulse generating regulator cells recover acute healing dynamics of some chronic conditions better than others. For example, in chronic conditions of low *ga*_1_ and low *ga*_3_, the sequential dynamics of each cell species are drastically improved. However, wound closure is not complete (Figures 5A and 5C). Alternatively, ideal wound closure, modeled as total cell number of species *C*_4_ becoming 100% of the cellular population in a timely manner, is observed in conditions with low *ga*_2_ and low *ga*_4_ signals production (Figures 5B and 5D).

**Figure 5:**
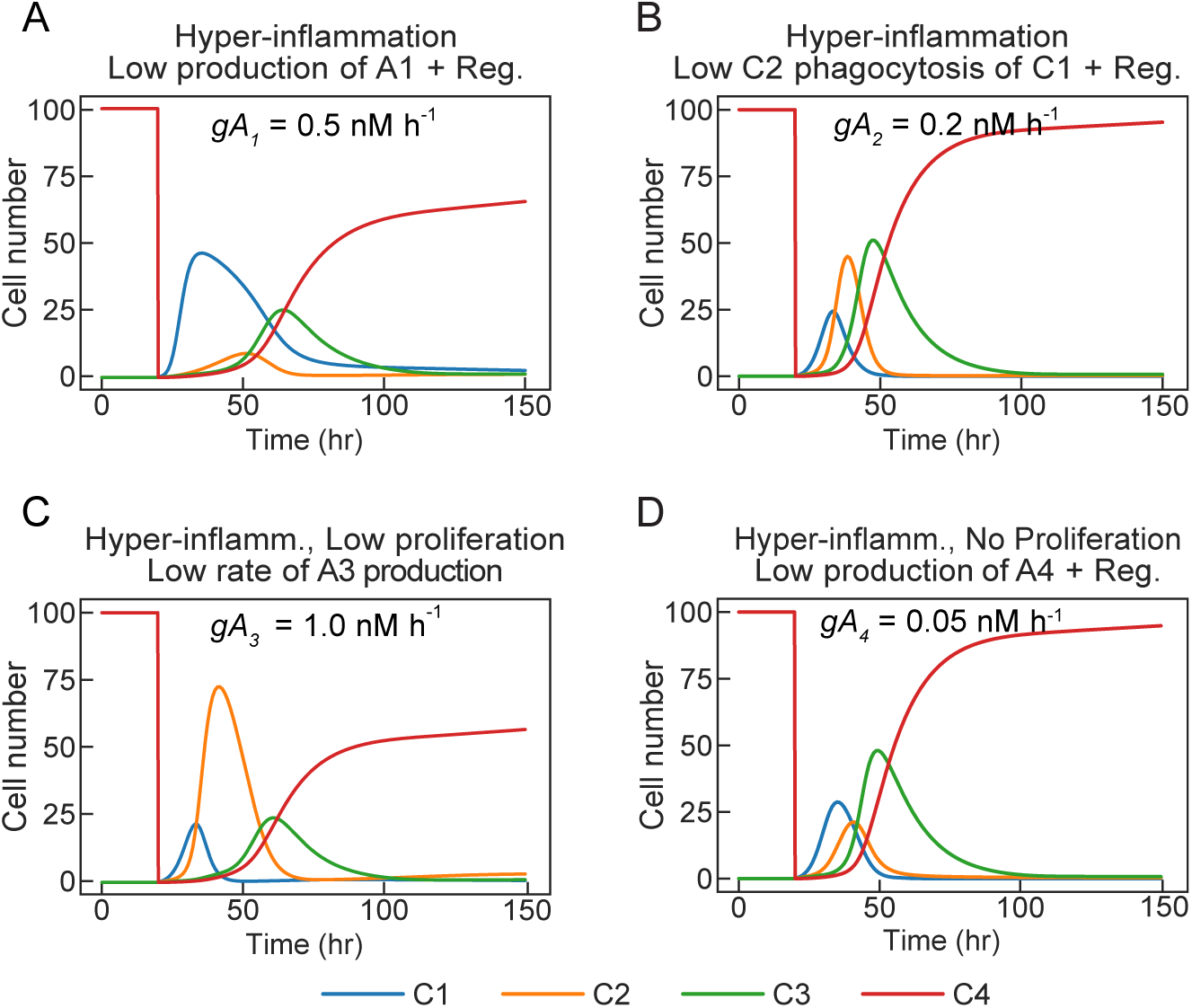
Simulations of chronic to acute wound healing dynamics with regulator cells. A. Low production rate of signal 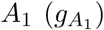, used to recruit *C*_2_. B. Low production rate of signal 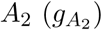, used to induce lysis expression in *C*_1_. C. Low production rate of signal 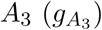, used to induce proliferation of cell *C*_3_ D. Low production rate of signal 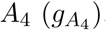, used to induce lysis expression in *C*_2_.

The regulator cells are most effective when signals *A*_1_ and *A*_3_ are abundant, but *A*_2_ and *A*_4_ are sparse. A low production rate of signal *A*_3_ proves to be difficult to regulate against chronic hyperinflammatory conditions. This is because *R*_2_ requires an abundance of *A*_3_ to activate the production of signal *A*_4_, to assist in healing progression. If signal *A*_3_ is absent (or low), the activation of the signal production of *A*_4_ in the regulator cell *R*_2_ is not effective in restoring healing dynamics (as shown in Figure 5C. However, if you compare the unregulated chronic dynamics (Figure 3C), to the dynamics of the regulated system, you will notice the healing dynamics improve due to decreased oscillatory dynamics of species *C*_1_, *C*_2_, and *C*_3_, although the final population of *C*_4_ does not change.

We also analyzed the stability of healing dynamics of the acute healing circuit layer by perturbing the growth rate, the death rate, and the signal production rate parameters of each cellular species in our circuit design. For example, in Figure 6, top row (i), we compare the system’s stability to *C*_1_ growth rate 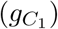, the production rate of signal 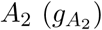, and *C*_1_ death rate 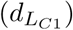 both in the absence (panel A) and presence (panel B) of regulator cells. Parameter perturbations were studied by either decreasing or increasing values listed parameter values in Table 2 as low as 0.1x and as high as 10x. Regulator cells are effective in increasing healing robustness when perturbing all signal synthesis parameters 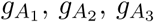, and 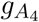. Contrarily, regulator cells struggle to improve healing dynamics of system sensitivity to cellular growth and death rates.

**Figure 6:**
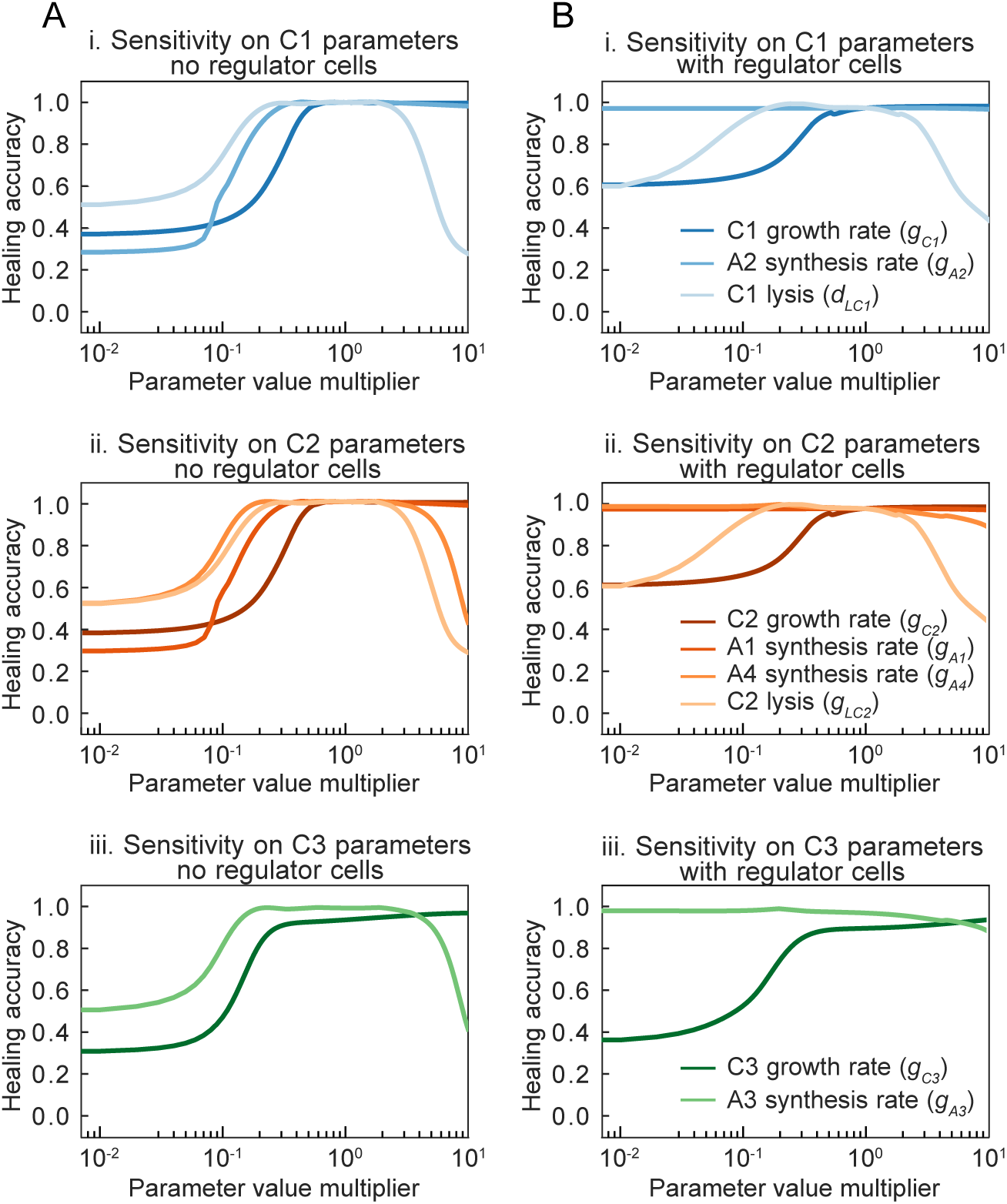
Analysis of the wound healing circuit dynamic stability to parameter perturbations. A. Simulations of circuit stability with parameter perturbations in the absence of regulator cells. B. Simulations of circuit stability with parameter perturbations in the presence of regulator cells. Parameter perturbations from 0.1 to 7.0 times ideal parameters listed in Table 2 acting on species (i.) *C*_1_, (ii.) *C*_2_, (iii.) *C*_3_. Healing accuracy is a measurement of complete and timely healing defined by the concentration of species *C*_4_ at time 150 hours. For example, 100% accuracy is species *C*_4_ reaching max concentration at time 150 hours.

## Discussion

Wound healing is a fundamental, yet complicated phenomenon developed to protect the host from intrusion of harmful pathogens. Chronic wound healing and effective treatments are important concerns for health care professionals. By taking advantage of new findings of signaling pathways and mechanistic relationships [25], we can better predict the stages of wound healing [26], develop anti-scarring therapies [27], better treat burn injuries, skin cancers, angiogenesis [28], and other chronic wound conditions [29].

Given the importance of signaling factors [30], there awaits new discoveries by coupling synthetic biology and wound healing signals for advanced treatments [31]. Computational studies on wound healing have provided the necessary insight into the molecular dynamics of wound healing phases, and the inter-connectedness of the complex signaling network. Researchers are currently applying classical control theory concepts to biological systems for sustainable mammalian-microbial interactions [32, 33, 34]. Implementing feedback controllers for regulation of immunological chronic diseases may prove to be a hands-off, stable solution for positively regulating complex network systems consisting of engineered multi-layered networks [35].

In this work we developed a simple model mimicking the cellular dynamics of wound healing, and designed biological controllers that can be embedded in a healing salve or placed on a bandage. Our computational approach focuses on the elucidation of control systems needed to sense and regulate against impaired dynamics of a sequential signaling network. We propose a multiple layer population controller consisting of a wound healing circuit that demonstrates cellular population dynamics of acute physiological wound healing (layer 1), and the coordination of feedback controllers that sense chronic dynamics, improving the system’s cellular dynamics to resemble acute healing (layer 2). In this paper we (1) simulate a four node wound healing process that resembles cellular dynamics found in physiological healing, (2) implement predictive chronic wound healing dynamics, and (3) regulate against hyper-inflammation and impaired proliferation chronic conditions using closed-loop controllers. Although we found conditions where our regulator controllers proved to be effective, there were still conditions in which our controllers failed to improve the chronic wound dynamics. x

We demonstrated the ability to use pulse generating motifs to sense temporal changes in chemical concentrations and cellular densities to predict and fix chronic conditions. We plan to continue to improve both the wound healing testbed layer, as well as develop more combinatorial network motif controllers to regulate against many more chronic conditions robustly. Having developed this model based on *E. coli* dynamics, we plan to build an experimental wound healing demonstration based on population controls and negative feedback motifs for predictable dynamics.

## Acknowledgments

The authors would like to thank Reed McCardell, William Poole, Ayush Pandey and Mark Prator for their insightful discussion. The authors L. G., X. R., and C. H. are supported by Defense Advanced Research Projects Agency (Agreement HR0011-17-2-0008). The content of the information does not necessarily reflect the position or the policy of the Government, and no official endorsement should be inferred.

## Notes

### Competing Interest Statement

The authors have declared no competing interest.

### Summary of Updates

Name change for a listed author.

